# Functional monocentricity with holocentric characteristics and chromosome-specific centromeres in a stick insect

**DOI:** 10.1101/2024.06.30.601385

**Authors:** William Toubiana, Zoé Dumas, Patrick Tran Van, Darren J. Parker, Vincent Mérel, Veit Schubert, Jean-Marc Aury, Lorène Bournonville, Corinne Cruaud, Andreas Houben, Benjamin Istace, Karine Labadie, Benjamin Noel, Tanja Schwander

## Abstract

Centromeres are specialized genomic regions that are critical for chromosome segregation in eukaryotes. However, how the diversity of centromeres evolved, ranging from localized centromeres in monocentric species to complex chromosome-wide distributed centromere regions in holocentric species, remains elusive. Our cellular and genomic analyses in *Timema* stick insects reveal that within cell variation of the major centromere protein CenH3 recapitulates the variation typically observed across species. While CenH3-containing nucleosomes are distributed in a monocentric fashion on autosomes and bind tandem repeat sequences specific to individual or small groups of chromosomes, they exhibit a holocentric-like distribution on the sex chromosome and bind to more complex intergenic regions. Despite this contrasting distribution, all chromosomes, including the sex chromosome, attach to spindle microtubules at a single location, marking the first instance of a functionally monocentric species with holocentric-like attributes. Together, our findings highlight the potential for gradual transitions towards holocentricity or CenH3-independent centromere determination, and help to understand the rapid centromere sequence divergence between species.

## INTRODUCTION

Chromosome segregation is a fundamental and conserved cellular process in eukaryotes, ensuring the transmission of genetic material to daughter cells during mitotic and meiotic cell divisions (1). It is governed by specialized chromosomal regions known as centromeres, where a unique histone H3 variant, referred to as CenH3 (or CenpA), is recruited to nucleosomes to replace the canonical H3 histone (2, 3). In turn, CenH3 facilitates the assembly of the kinetochore protein complex, which mediates the attachment of spindle microtubules (4-6). In the majority of eukaryotes, CenH3 binds to DNA sequences at a single, well-defined centromere region on each chromosome (7). Centromere sequences in these so-called monocentric species are typically composed of tandem or interspaced repeats which are absent or rare in non-centromeric regions of the genome (7). By contrast, multiple lineages have independently evolved a holocentric chromosome structure, where CenH3 proteins and microtubules attach along the entire length of all chromosomes (8, 9). In these lineages, centromere sequences vary, and can comprise repeated DNA motifs akin to those in monocentric species, or more complex DNA structures forming broad intergenic or poorly transcribed domains (10-14).

Irrespective of the variation that exists between species, chromosomes within a species always share the same centromere configuration (e.g., holo- or monocentric), the same centromere composition (e.g., tandem repeats, transposons or complex DNA structures) and typically similar CenH3 binding sites (e.g., similar tandem repeat or transposon sequences) (10, 12, 13, 15-18). This uniformity in centromere structure within species has notably led to the argument that diversity in centromere composition and configuration arises from discrete evolutionary transitions, without intermediate states (19).

While investigating the evolution of centromeres within the stick insect genus *Timema*, we uncovered a novel spatial organization and recruitment dynamic of the CenH3 protein, involving variations that are reminiscent of both mono- and holocentric species. We substantiated our intriguing cytological observations with chromatin immunoprecipitation and sequencing (ChIP-seq) data, which further revealed that CenH3 binds to strikingly different sequences, encompassing both tandem repeats and more complex DNA structures. Finally, we also observed unexpected variability in the role of CenH3 to assemble kinetochores and recruit spindle microtubules, thus questioning its primary role in defining centromeres. Together, our findings indicate a possible intermediate state in transitions from mono-to holocentricity, or towards a CenH3-independent centromere definition.

## RESULTS

### A CenH3 distribution reminiscent of holo- and monocentric species

*Timema* is a genus of stick insects with multiple independent transitions to asexuality (female-producing parthenogenesis) (20). This context provides a unique opportunity to investigate the evolution of centromeres under different selective conditions. While characterizing centromeres in different *Timema* species, we conducted immunostaining on male gonads of *T. douglasi* using a custom antibody for the CenH3 protein (Figure 1). As CenH3 plays a central role in centromere identity and chromosome segregation in eukaryotes, our focus was directed towards cells in meiotic metaphase, when spindle microtubules bind to centromeres before segregation initiates (4).

**Figure 1:**
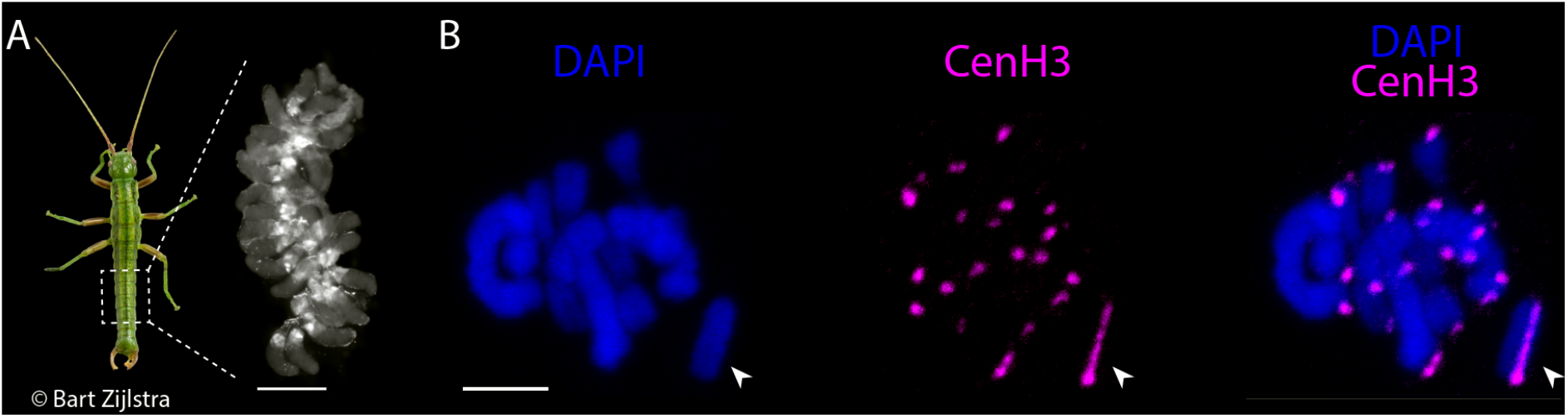
Differential CenH3 binding between autosomes and the X chromosome at metaphase I during male meiosis. A) Image of a *Timema* male and a gonad containing meiotic and spermatid cells at various stages of maturation. The dashed square highlights the abdominal region housing the pair of gonads. Scale bar corresponds to 400 micrometers. (B) Ring and rod bivalents of paired homologous autosomes in metaphase I. The univalent sex chromosome (X chromosome) is indicated by the arrowheads. At this stage, CenH3 exhibits a monocentric distribution on the autosomes and a holocentric-like distribution on the X chromosome. Scale bar corresponds to 5 micrometers.

During metaphase I, we observed striking differences in CenH3 distribution between the autosomes and the X chromosome. On each of the 22 autosomes, CenH3 formed a single, distinct focus (Figure 1B). This pattern is characteristic of species with monocentric chromosomes and was expected for *Timema* given their karyotypes with chromosomes featuring clear primary constrictions (21). Conversely, the univalent X chromosome exhibited binding of CenH3 along its entire length, akin to species with holocentric chromosomes (Figure 1B). Note that *Timema* have an X0 sex chromosome system, meaning that there is a single X and no Y chromosome in males (21). We confirmed the different CenH3 binding patterns between the X and autosomes in a CenH3-directed ChIP-seq assay on male gonads, where we mapped ChIP and input derived sequence reads to a newly generated chromosome-level genome assembly of the same species (SAMN41832294; see Material and Methods; Supplemental figure 1; Supplemental table 1). While each of the autosomes displayed a distinct, monocentromere-typical ChIP signal, the X chromosome was characterized by an enhanced signal distributed over its entire length, with a somewhat increased intensity at one chromosome end (Figure 2A).

**Figure 2:**
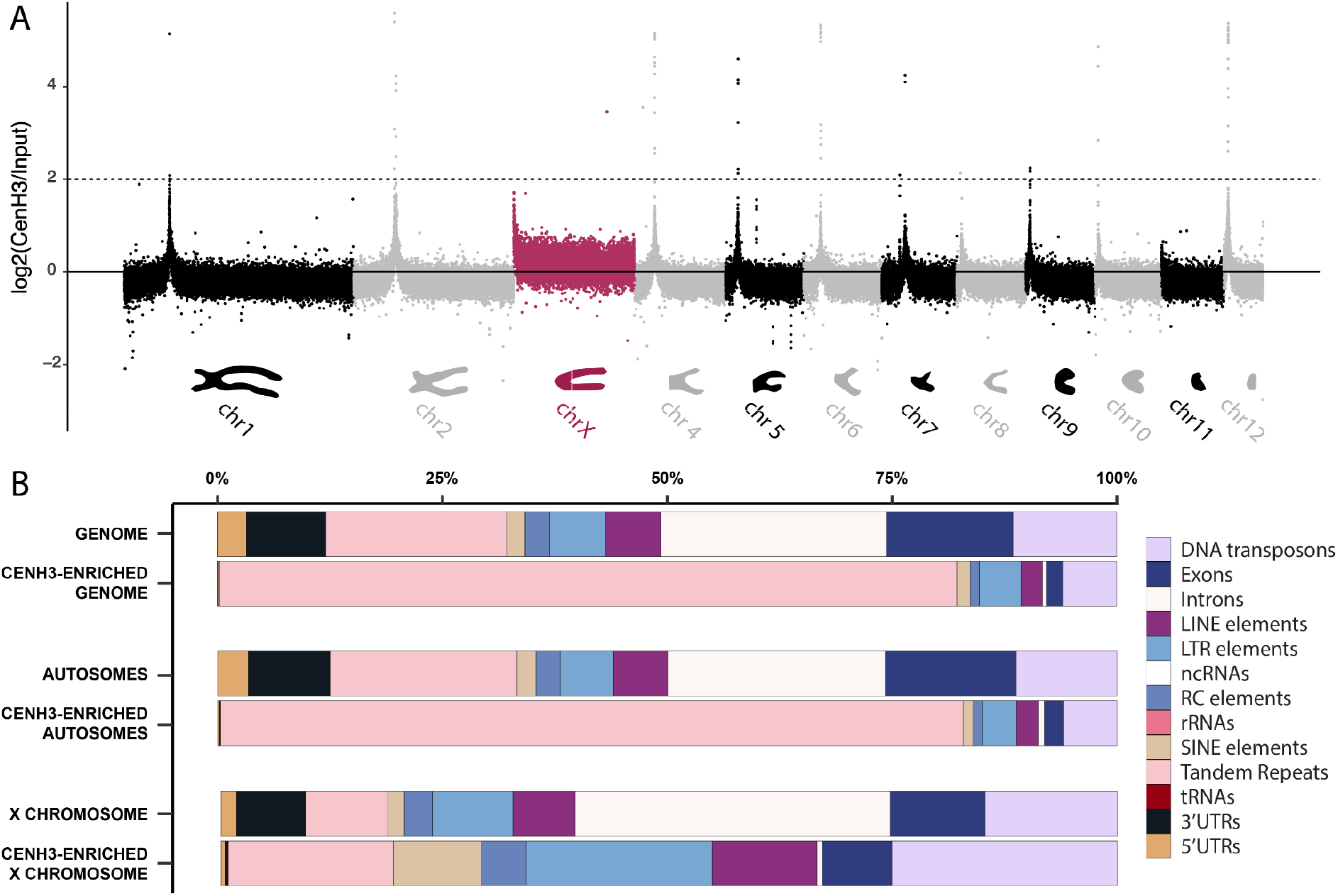
CenH3-ChIP signal corroborates differences in CenH3 distribution between autosomes and the X chromosome. A) Depiction of the CenH3 directed ChIP-signal in non-overlapping 10kb windows (log2 transformed coverage ratio of CenH3 immunoprecipitation derived reads versus control reads without immunoprecipitation (input) along each chromosome (x-axis). The schematic representation of the chromosome morphologies (modified from (21)) indicates the approximate locations of the primary constrictions. The dashed line denotes the threshold used to define CenH3-enriched regions (i.e., centromere windows, see Material and Methods). B) Representation of thirteen genomic features observed genome-wide, on autosomes, and the X chromosome, and enriched in CenH3-directed ChIP.

Given the highly unexpected CenH3 distribution on the X chromosome of *T. douglasi* males, we investigated whether it was specific to the focal species or a general feature in the *Timema* genus. We used males from three additional species, which cover the phylogenetic breadth of the genus, to conduct CenH3 staining as described for *T. douglasi*. Our cytological observations were consistent across all species (Supplemental figure 2), indicating that the different CenH3 distribution between autosomes and the X chromosome is a conserved feature in *Timema* males.

### The holocentric-like CenH3 recruitment is dynamic and extends beyond the X chromosome in early male meiosis

To elucidate the cellular mechanism(s) governing CenH3 distribution along the X chromosome, we examined CenH3 localizations at earlier meiotic stages. We assigned individual cells to specific stages using the Structural Maintenance of Chromosomes protein 3 (SMC3), a subunit of the cohesin complex that aids in recognizing key steps during meiosis (22). Surprisingly, we found that at the onset of meiosis, the holocentric CenH3 distribution is not restricted to the X chromosome. Instead, CenH3 is recruited along the entire length of all chromosomes, colocalizing with SMC3 (Figure 3). This holocentric-like CenH3 recruitment was most prominent during the leptotene and zygotene stages of prophase I when sister chromatid cohesion was established, and homologous chromosome synapsis was only initiated (Figure 3). As synapsis progressed, CenH3 became gradually condensed into a discrete focus on each autosome (Figure 3, zygotene and pachytene stages), while its holocentric-like distribution persisted on the univalent X chromosome (Figure 3, pachytene and metaphase I stages). Finally, at the onset of meiosis II, when sister chromatid arms are no longer attached together and the cohesin complex is removed from chromosome axes, the holocentric-like CenH3 distribution on the X disappeared, such that a single focus was visible on all chromosomes (Figure 3, prophase II stage). In summary, the entry of meiosis, generally associated with cohesion of sister chromatids and chromatin compaction via loop extrusion (23), is characterized by CenH3 recruitment along the entire length of all chromosomes in *Timema*. Once synapsis is completed, only the X chromosome maintains a holocentric CenH3 distribution, most likely because it does not have a homologous partner to synapse with in males.

**Figure 3:**
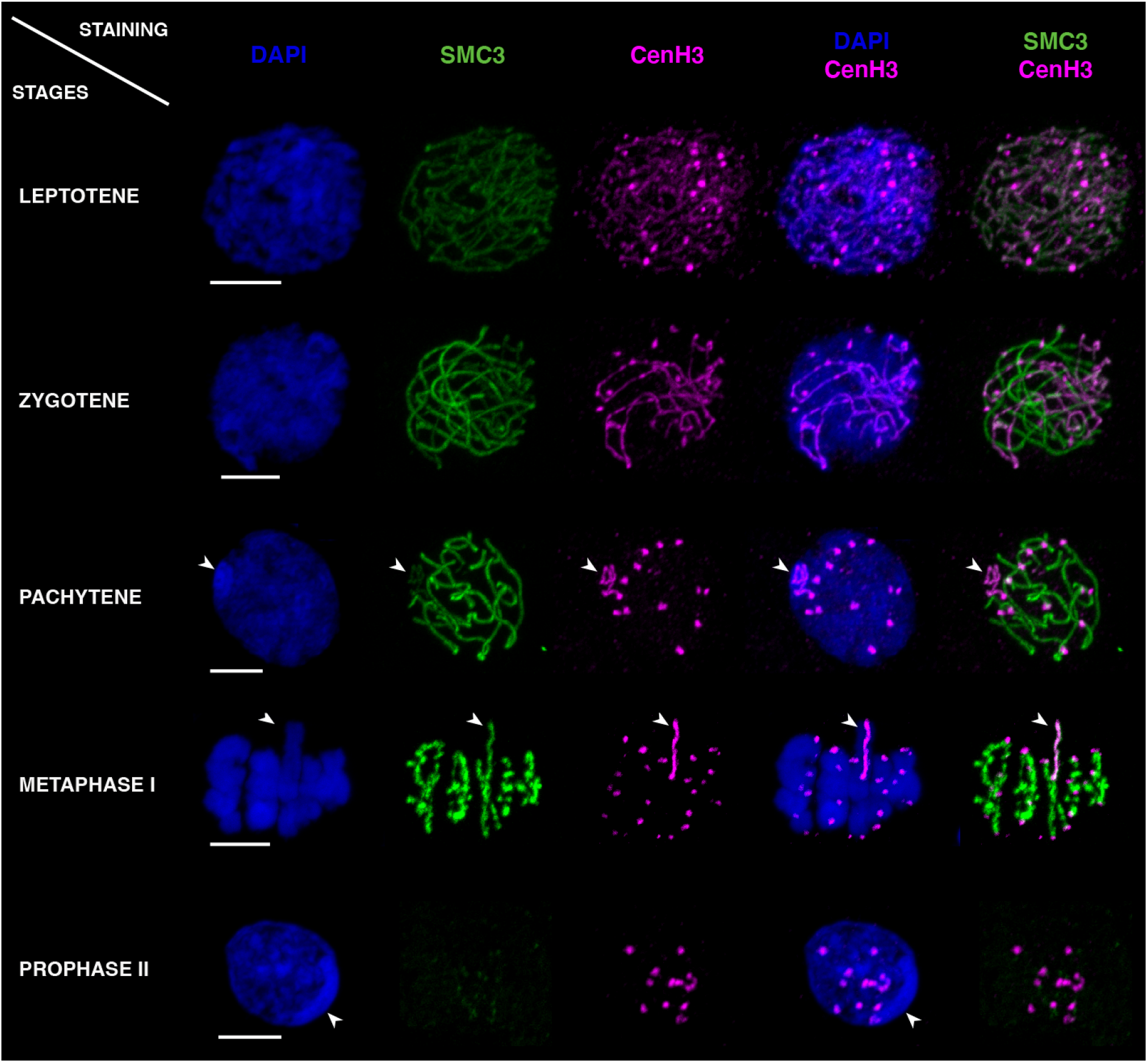
Dynamics of CenH3 and SMC3 distributions along *T. douglasi* chromosomes during prophase I and II. The labeling highlights DNA with DAPI staining in blue, SMC3 in green, and CenH3 in magenta. During the leptotene and zygotene stages, CenH3 and SMC3 co-localize along the entire length of all chromosomes. CenH3 then retracts to single foci on all chromosomes except the X during pachytene, while SMC3 remains distributed along the entire length of all chromosomes until metaphase I. Prophase II is marked by the disappearance of the holocentric-like distribution of CenH3 on the X chromosome and SMC3 retracting to single foci near the monocentrically distributed CenH3. When discernible, the X chromosome is indicated by arrowheads. Scale bars correspond to 5 micrometers.

### CenH3 binding sites differ between the X and autosomes and reveal distinct autosomal centromeres

We then investigated whether the distinct CenH3 distributions on autosomes *versus* the sex chromosome were mirrored by distinct CenH3 binding sites. To this end, we used our CenH3-directed ChIP-data on male gonads and first tested for enrichment of specific sequence categories (i.e., exons, introns, tandem repeats, DNA transposons, etc., Figure 2B). Enrichment patterns differed strikingly between the autosomes and the X. For the autosomes, the CenH3-ChIP data were predominantly composed of tandem repeat sequences, with a highly significant enrichment relative to the genomic background (Figure 2B; chi-square = 1941157, DF= 12, p < 0.001; see Material and Methods). For the X chromosome, no specific sequence category was overrepresented (Figure 2B). Instead, the CenH3-ChIP data had a reduced representation in genic regions (i.e., introns, exons and UTRs), which largely drove the significant difference in sequence categories relative to the genomic background (Figure 2B; chi-square = 189028, DF= 12, p < 0.001).

We further used our CenH3-ChIP data to characterize centromere sequences in *T. douglasi*. Given the enrichment in tandem repeats in the CenH3-ChIP data, we identified specific tandem repeat families constituting centromeres by using an assembly-based and assembly-free (i.e., k-mer based) approach (see Material and Methods; Supplemental figure 3). For both approaches, individual repeats were grouped into families using network clustering analysis which resulted in 28 and 16 centromere repeat families, respectively, with closely matching motif sequences (Supplemental figures 3A, 3C; Supplemental tables 2, 3). Among these, six dominant repeat families (repeat family 1, 2, 7, 10, 13 and 22; Supplemental table 2) collectively constituted more than 90% of the tandem repeats found in centromeric regions. One of these abundant families (repeat family 2) comprised the AACCT motif, a telomere repeat in various insect lineages (Figure 4A; Supplemental figures 3, 4; Supplemental table 2; (24, 25)). Four other families (repeat families 1, 7, 13 and 22) consisted of motifs ranging from 63 to 384 base pairs in size. Finally, the sixth family (repeat family 10), with a motif size of ∼1400 bp, was a composite of families 2 and 7 (Supplemental figures 3, 4; Supplemental table 2) and was therefore not considered further.

**Figure 4:**
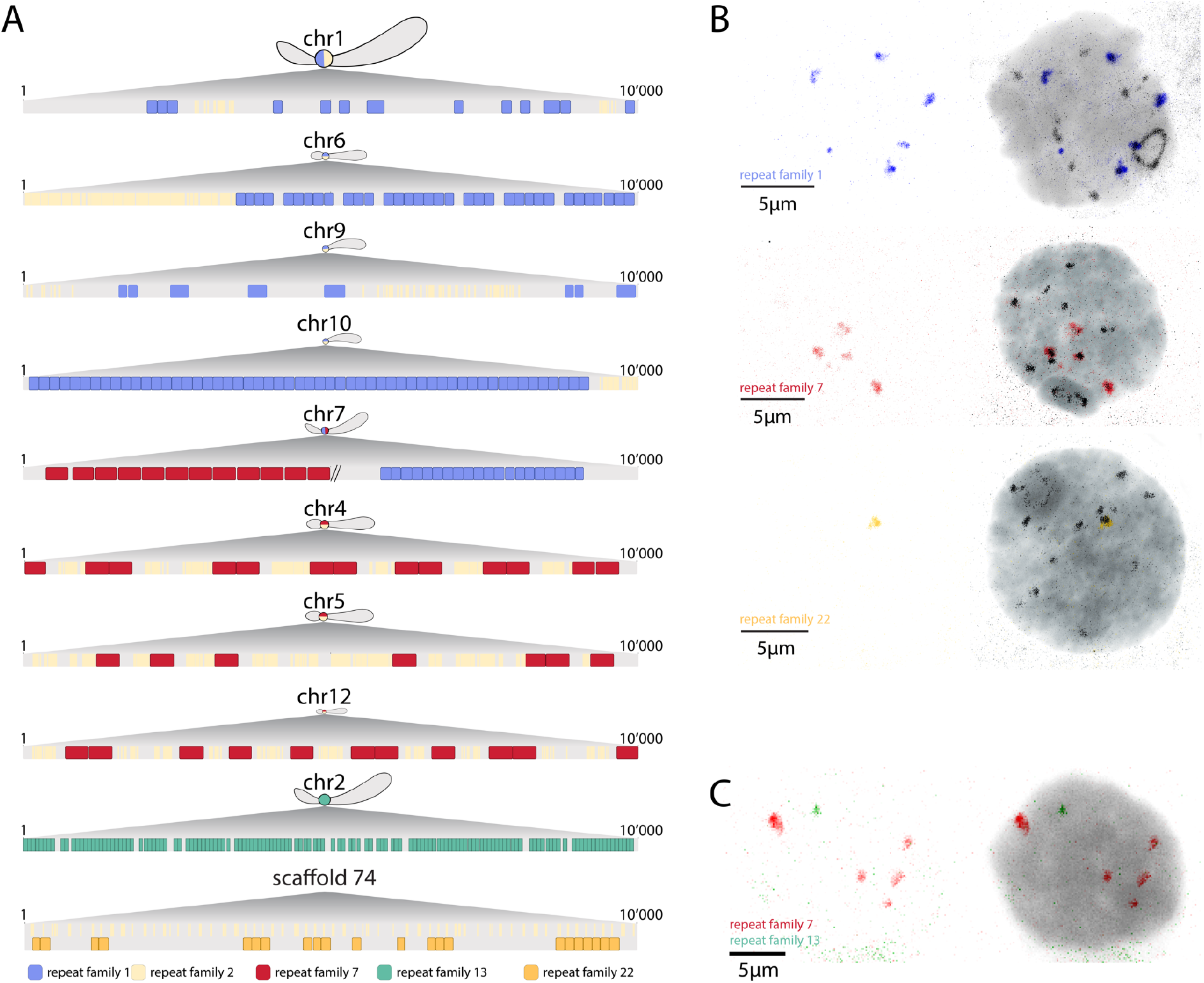
The centromere sequence composition differs between *T. douglasi* autosomes. A) 10kb regions with the most enriched CenH3 ChIP signal, color-coded with the five most abundant centromere repeat families representing over 90% of the total tandem repeat content. Chromosome drawings indicate the centromere localization within each autosome, including sub-metacentric (chromosomes 1, 2 and 7), acrocentric (chromosomes 4, 5, 6 and 12) and telocentric (or near telomeres; chromosomes 9 and 10) chromosomes. Note that the telocentric chromosomes 8 and 11 are missing because of incompletely assembled centromeres (see text). B) Immuno-FISH on meiotic cells labeled with CenH3 proteins (black) and DNA probes targeting different centromere repeat families. C) FISH staining highlighting the location of two centromere repeat families, revealing that they are on distinct chromosomes (non-overlapping).

We examined the representation of the five distinct repeat families (repeat family 1, 2, 7, 13 and 22) in 9 out of the 11 autosomal centromeres. The incomplete assembly of centromeres of the two remaining autosomes (chromosomes 8 and 11), both telocentric (Figure 2A), precluded their in-depth examination. The distribution of the five repeat families among the nine more complete autosomes revealed significant differences in centromere composition (Supplemental figures 4A, 4B; Supplemental table 4). In four out of these nine autosomes (chromosomes 1, 6, 9, 10), centromeres consisted of two repeat families (family 1 and 2) which were each tandemly repeated (Figure 4A). The centromeres of three additional autosomes (chromosomes 4, 5, 12) consisted of two repeat families (family 2 and 7) organized into higher-order repeats (Figure 4A). One autosome (chromosome 7), contained repeats present in the centromeres of two otherwise distinct groups of autosomes (Figure 4A) while chromosome 2 had a unique, chromosome-specific centromere sequence (Figure 4A). Overall, repeat family 2 (corresponding to the telomeric AACCT motif) was the most broadly shared, as it was found in high abundance in the centromeres of seven autosomes (chromosomes 1, 4, 5, 6, 9, 10 and 12).

We corroborated the inferred centromere sequences for individual or groups of autosomes through a combination of Fluorescence In Situ Hybridization (FISH) and immuno-FISH experiments. Thus, repeat family 7, which was predicted to be present in the centromeres of four autosomes (chromosome 4, 5, 7, and 12), labeled four centromeres along with CenH3 (Figure 4B). Repeat family 13, which was predicted to be specific to chromosome 2, labeled a single chromosome (Figure 4C). Repeat family 22, which was not represented in the centromeres of the nine more complete autosomes but only on four unanchored scaffolds (Figures 4A; Supplemental figures 4A, 4B), also labeled a single chromosome (Figure 4B; Supplemental figure 5). Finally, repeat family 1 was predicted to be present on five autosomes and indeed labeled five autosomes as well as one extremity of the X chromosome (Figures 4A, 4B). Importantly, each of the examined repeat families exhibited distinct nuclear locations, indicating that they are in centromeres of different chromosomes (Figure 4C; Supplemental figure 5). Collectively, these findings highlight the divergence of centromere sequences among different autosomes of *T. douglasi*, which mirrors the well documented rapid divergence of centromere sequences between species (7, 19, 26). The reasons why some chromosomes have unique (chromosome-specific) centromere sequences while others form groups of similar centromere sequences remains to be investigated. Admixture between divergent populations or species could result in centromere heterogeneity, yet introgression analyses in *Timema* do not point to hybrid origins of *T. douglasi* (20). Another possible situation favoring centromere sequence convergence among specific sets of chromosomes is nuclear compartmentalization, where certain chromosomes or certain centromeres cluster into specific regions of the nucleus (27, 28). Such spatial clustering could facilitate convergent evolution between interacting centromere repeats on different chromosomes.

**Figure 5:**
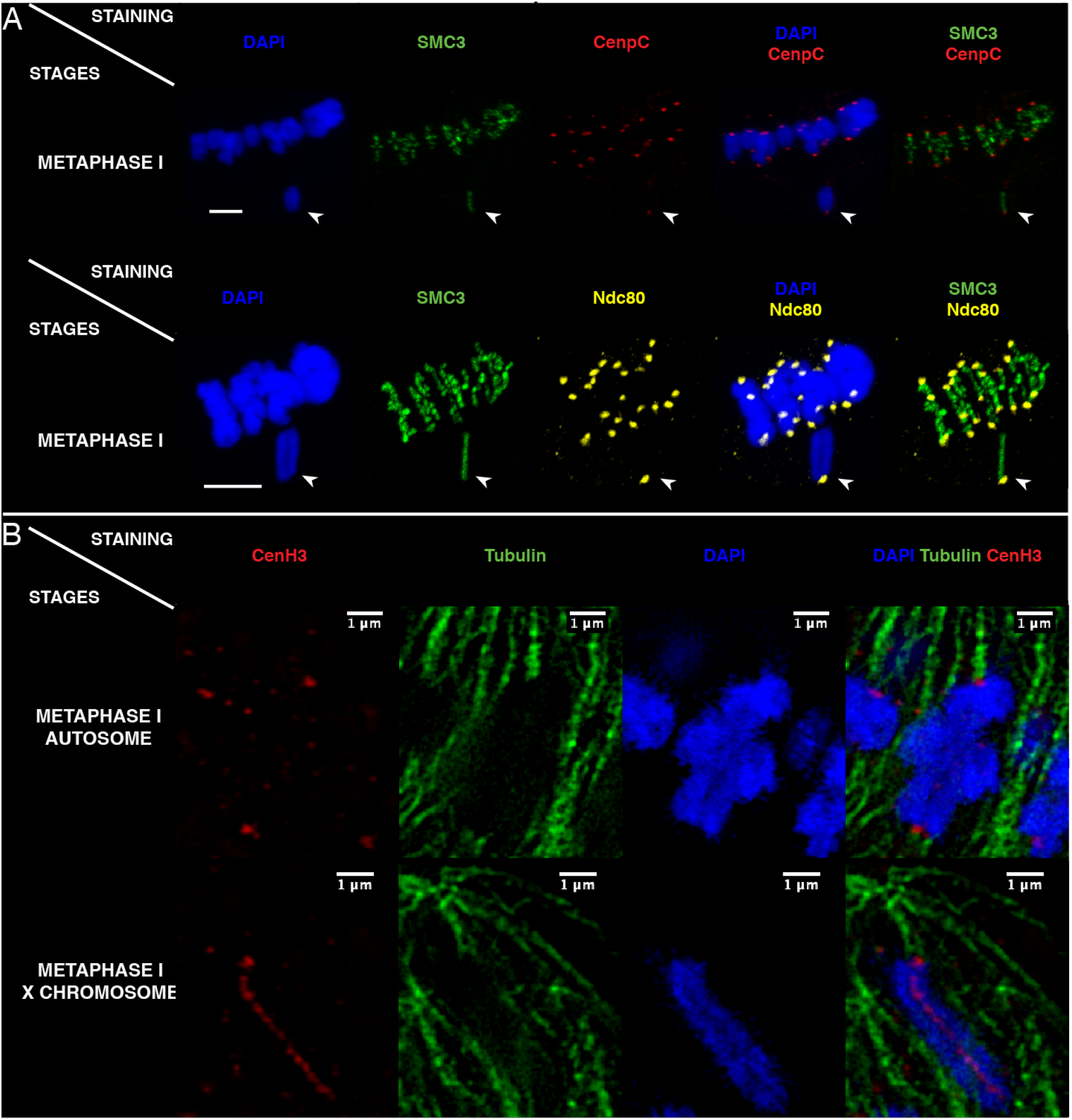
Kinetochore and spindle microtubule distributions on *T. douglasi* metaphase I chromosomes. A) All chromosomes, including the X chromosome (arrowheads), showed monocentric-typical CenpC (red) and Ndc80 (yellow) signals at the poleward chromosome termini. Scale bars correspond to 5 micrometers. B) 3D-SIM (single slice of an image stack) visualized the monocentric microtubule attachment sites along with CenH3 on autosomes and the X chromosome during metaphase I.

Regarding the X chromosome, our immuno-FISH experiments indicated that in addition to the holocentric CenH3 distribution, one chromosome end likely harbors a localized centromere, composed of family 1 repeats (Figure 4B). This chromosomal region was also distinctly visible as a somewhat increased ChIP signal on the left end of the X assembly (Figure 2A), while featuring an enriched CenH3 immuno-labeling signal (Figures 1B, 4B) and matching the telocentric phenotype of the X chromosome.

### Kinetochore protein distributions and microtubule attachments suggest a functionally monocentric X chromosome

To determine whether the holocentric-like CenH3 distribution translates into a functionally holocentric X chromosome during segregation, we investigated the distribution of kinetochore proteins and utilized super-resolution 3D-SIM to localize microtubule attachment sites. As expected for a functionally monocentric X chromosome, CenpC and Ndc80 proteins (members of the inner and outer kinetochore region, respectively) were distinctly visible at a unique position on the X at metaphase I, resembling the CenH3-monocentric autosomes (Figure 5A). Similarly, alpha-tubulin and CenH3 co-staining analyzed using 3D-SIM ((29); see Material and Methods), revealed monocentric spindle attachments to a delimited and terminal region of the X chromosome as well as autosomes (Figure 5B). Together, these findings indicate that despite the holocentric-like distribution of CenH3, the X chromosome behaves as a functionally monocentric chromosome during the first meiotic division in *Timema* males.

To our knowledge, this study represents the first description of a lack of co-occurrence of CenH3 with certain kinetochore and spindle microtubule proteins in animals and monocentric species more generally. In addition to suggesting two distinct recruitment steps for centromere proteins, our observations contrast the findings in human cell lines and *Drosophila* mutants where the mis-incorporation of CenH3 leads to an ectopic assembly of partial or full kinetochore complexes (30, 31).

## DISCUSSION

Centromeres and variation in centromere organization have been known since early work on karyotype evolution in plants and animals (9), yet our understanding of how these key structures evolve and diversify remains limited (19). For example, how transitions from monocentric to holocentric centromere configurations can occur over the course of evolution, and whether these transitions are sudden or gradual is unknown (19). In *Timema*, the centromere histone variant CenH3 can show a holocentric or monocentric distribution within the same cell, with the distribution varying between chromosome types and meiotic stages. This plasticity in CenH3 binding highlights the potential for rapid evolution between different centromere configurations such that it may elucidate how holocentric transitions have repetitively occurred across eukaryotes (9).

Our study also illustrates that transitions to holocentricity can evolve gradually. For example, CenH3 could acquire the ability to bind along chromosome lengths before entailing functional holocentricity. Indeed, our findings reveal that the holocentric distribution of CenH3 can be decoupled from the holocentric recruitment of kinetochore proteins and holocentric segregation of chromosomes. The disparate distributions of CenH3 and kinetochore proteins may stem from the distribution of the CenH3 proteins facing inwards the two chromatids. Holocentric lineages in plants and animals typically have their CenH3-enriched chromatin facing outwards the two chromatids such that two linear lines form on every chromosome (8, 10, 14, 32, 33). By contrast, the *Timema* X chromosome exhibits a single CenH3 line positioned in between the two chromatids, potentially impeding the assembly of a functional kinetochore and subsequent holocentric segregation. Alternatively, the disparate distributions between CenH3 and kinetochore proteins in *Timema* may represent a first step towards a CenH3-independent assembly of the kinetochore complex. CenH3-independent centromeres have repeatedly evolved within insect phylogeny, in association with transitions to holocentricity (34, 35). Under this scenario, *Timema* would represent an intermediate state between CenH3-independent holocentric lineages and CenH3-dependent monocentric lineages. More generally, the fact that CenH3 alone is not sufficient to drive kinetochore assembly and microtubule recruitment in *Timema*, as well as in some holocentric insect and plant lineages (35-37), questions the view that CenH3 is the key epigenetic “mark” defining centromeres.

Another knowledge gap in centromere evolution is to understand how centromere sequences can evolve so rapidly among species as this should require parallel rapid evolution of DNA-centromere protein affinities (19, 26). Indeed, in most studies where centromere sequences have been formally characterized, comparisons point towards high inter-species variation but low divergence between chromosomes within species (7, 10, 18, 38). Our study contributes to filling this knowledge gap by revealing that CenH3 binds to divergent centromere sequences on different autosomes, similar to what has been recently documented within specific lineages (39-42). Moreover, CenH3 in *Timema* can bind to broad categories of intergenic regions, as revealed on the X chromosome, mirroring the centromere domains of several holocentric lineages (12-14). Such versatility in binding affinity within the same cell could facilitate a rapid turnover of centromere sequences between species, including those associated with transitions to holocentricity.

Overall, *Timema* offers a new perspective on the evolution of centromeres, which calls for an increased phylogenetic diversity in model organisms used to study key cellular features. The fact that chromosomes generally adopt either a monocentric or holocentric configuration within a lineage has been associated with the idea that centromere complexity evolves through discrete transitions (19). Our cellular and molecular observations suggest that gradual transitions towards holocentricity are possible, and may involve intermediate states that can remain stable over long evolutionary timescales. Finally, the turnover in centromere sequence is hypothesized to result from centromere competition during asymmetric female meiosis (26). Because we identify striking centromere sequence divergence between non-homologous chromosomes that do not compete during meiosis, our findings raise the question of other drivers of sequence divergence and whether cellular processes such as chromosome interactions also contribute to centromere sequence diversity.

## Supporting information

Supplemental material and results

## ACKNOWLEDGEMENTS

We thank Maria Cristina Gambetta, Corinne Peter and the Lausanne GTF platform for help with ChIP-seq, Severine Lorrain from PAF for help with Western Blot, Bart Zijlstra for the *Timema* picture in Figure 1, Virginie Hamel and Paul Guichard for help with super-resolution microscopy, Marie Delattre, Alexander Woglar, Pierre Gönczy for discussions, Qiaowei (Miya) Pan and Hugo Darras for discussions and comments on the manuscript, and current and previous members of the Schwander lab for discussions.

## Author contributions

W.T. and T.S. designed the study. W.T. and Z.D., C.C. and K.L. performed molecular work. W.T., Z.D., L.B., J-M.A., B.I. and B.N. developed methods. W.T., D.J.P., P.T.V., V.M. and V.S. analyzed the data with input from T.S. and A.H., and W.T. and T.S. wrote the paper with input from all authors.

## Data and code availability

Raw sequence reads used for the reference genome have been deposited in NCBI’s sequence read archive under bioproject PRJEB75180. Data were processed to generate plots and statistics using R v3.4.4.

