## Supplemental material and results for "Functional monocentricity with holocentric characteristics and chromosome-specific centromeres in a stick insect"

### Supplemental Materials

### Methods

#### Sample collection and reference genome generation

We used wild-collected individuals of the species *T. douglasi* (individuals for CenH3 related questions: 38°57'24.9"N 123°32'10.0"W; individuals for the reference genome: 38°58'56.4"N 123°28'11.9"W), *T. knulli* (35°50'10.3"N 121°23'29.3"W), *T. californicum* (37°20'35.4"N 121°38'11.3"W), and *T. bartmani* (34°09'48.1"N 116°51'43.5"W). To generate the *T. douglasi* reference genome, we assembled contigs based on Nanopore and Illumina libraries (sequenced at 62x and 63x coverage respectively) generated from a single female, and then scaffolded contigs using a Hi-C library (sequenced at 95x coverage) based on a different female. We annotated this assembly using transcriptome data from different tissues and development stages of males and females of *T. douglasi* as well as from a closely related species (*T. poppense*; see Gene annotation section).

##### Sequencing libraries

To extract high molecular weight (HMW) DNA we flash-froze a single female (without gut) in liquid nitrogen and ground it using a Cryomill (Retsch). We then extracted HMW DNA using a G/20 Genomic Tips kit (Qiagen) following manufacturer's protocols. We checked DNA integrity on a pulse field agarose gel.

A total of 4 ONT libraries were prepared following Oxford Nanopore instructions. One library was prepared using the SQK-LSK108 ligation sequencing kit and was loaded on a MinION R9.4.1 Flow Cell, and three libraries were prepared using the SQK-LSK109 ligation sequencing kit and were loaded on PromethION R9.4.1 Flow Cells. Flow Cell loading was performed according to the Oxford Nanopore protocol and resulted in 62X coverage.

A PCR free Illumina library was prepared using the Kapa Hyper Prep Kit (Roche, Basel, Switzerland), following the manufacturer's instructions. Library was quantified by qPCR using the KAPA Library Quantification Kit for Illumina Libraries (Roche), and library profile was assessed using a High Sensitivity DNA kit on an Agilent Bioanalyzer (Agilent Technologies, Santa Clara, CA, USA). The library was then sequenced to approximately 66x coverage on an Illumina HiSeq 4000 instrument (Illumina, San Diego, CA, USA), using 150 base-length read chemistry in a paired-end mode.

Hi-C library construction using the Proximo Hi-C Kit and sequencing (250 Mio read pairs) was outsourced to Phase Genomics (Seattle). We generated ground tissue for cross-linking following the manufacturer's protocol, using a different female than the one used for Nanopore and NovaSeq sequencing, from the same natural population.

#### Assembly pipeline and parameters

Raw Oxford Nanopore reads were filtered using Filtlong v0.2.0 ([https://github.com/rrwick/Filtlong](https://github.com/rrwick/Filtlong)) with the parameters --min_length 1000 --keep_percent 90 --target_bases 69050000000. The filtered Nanopore reads were then assembled into contigs using Flye v2.8.1 (1) with --genome-size 1.3 Gbp. All Nanopore reads were mapped against the contigs using minimap2 v2.19 (2) with the parameters -c -x map-ont and a first step of polishing was performed using Racon v1.4.3 (3). Three additional rounds of polishing were then conducted using the Illumina short reads. The short reads were aligned to the contigs using BWA mem v0.7.17 (4) and polishing was performed using Pilon v1.23 (5).

The assembly was decontaminated using BlobTools v1.0 (6) under the taxrule "bestsumorder". Hit files were generated after a blastn v2.10.1+ against the NBCI nt database, searching for hits with an e-value below 1e-25 (parameters: -max_target_seqs 10 -max_hsps 1 -evalue 1e-25). Contigs without hits to metazoans were removed. Haplotypic duplications were filtered out: filtered reads were mapped against the decontaminated genome using minimap2 and haplotigs were detected with Purge Haplotigs v1.1.1 (7) using the parameters -l 3 -m 17 -h 190 -j 101 following the recommendations by (7).

For scaffolding, Hi-C reads were mapped to the haploid genome using Juicer v1.6 (8) with the restriction site Sau3AI. Chromosome-level scaffolding was then performed using 3D-DNA v180922 (9) with the parameters --editor-coarse-resolution 25000, as recommended by the authors. The resulting Hi-C contact matrices were visualized with Juicebox, and polished following the recommendations by (8). The completeness of the assembly was assessed with BUSCO v5.1.2 (10) and the *insecta_odb10* dataset using the --long and --augustus parameters.

To identify the X chromosome in our assembly, we used a coverage approach. We compared coverage between males and females because *Timema* have XX/X0 sex determination (11) and males are expected to show half of the female coverage at the X chromosome. We mapped 5 female (SRS7637469, SRS7637489, SRS7637497, SRS7638280, SRS7638278 from (12)) and 2 male samples (Bioproject PRJNA808673 from (13)) to our scaffolded genome which allowed us to unambiguously identify the third largest scaffold as the X chromosome (Supplemental figure 1).

#### Genome annotation

##### Gene annotation

The *T. douglasi* genome was annotated using a combination of *ab initio* gene prediction, protein homology, and RNA-seq using the Braker2 pipeline v. 2.1.6 (14). To begin, the genome assembly was soft-masked using RepeatModeler (v. 2.0.2, options: -LTRStruct, -engine ncbi) and RepeatMasker (v. 4.1.2, options: -engine ncbi, -xsmall). For protein evidence, we used the arthropod protein sequences from OrthoDB v.10.1 (15) and the predicted protein sequences for *Timema* from our previous genome assemblies (12). For RNAseq evidence, we used publicly available RNAseq data from *T. douglasi* and *T. poppense* (Bioproject Accessions: PRJNA380865, PRJNA1128519, and PRJNA1128554). This is a total of 376 RNAseq libraries (364 paired-end, 12 single-end) covering 117 different life stages, tissue, and sex combinations. Reads were quality trimmed with Trimmomatic (v. 0.39, options: ILLUMINACLIP:3:25:6 LEADING:9 TRAILING:9 SLIDINGWINDOW:4:15 MINLEN:80) (16) before mapping to the genome assembly with STAR (v. 2.7.8a, options: --twopassMode Basic) (17). Braker2 was run using protein evidence and RNAseq separately with the gene predictors Augustus v. 3.4.0 (18) and Genemark v. 4.72 (19). Following the RNAseq run, UTR predictions were added to the RNAseq gene predictions using GUSHR v. 1.0 (20) in Braker2 (--addUTR=on). The separate gene predictions were then merged using TSEBRA v1.0.3 (21) using the pref_braker1.cfg configuration file, which weights RNAseq evidence more strongly than the default option. We then ran BUSCO (v. 5.3.2, insecta_odb10) on the gene regions annotated by Braker2 and on the whole genome assembly. Any genes found by BUSCO but missed by Braker were then added to the annotation (48 genes). ncRNA genes were predicted using Infernal (v. 1.1.2, minimum e-value 1e-10) (22). GO-terms for protein-coding gene predictions were obtained using blastP within OmicsBox v3.1.2, default parameters) to blast the nr *Drosophila melanogaster* database (taxonomy filter: 7227).

##### Transposable element and tandem repeat annotations

Transposable elements were annotated using RepeatModeler2 v. 2.0.3 (23). A library of consensus sequences of repeats was built and annotated using eleven *Timema* assemblies. Structural detection of LTR elements was activated using the -LTRStruct option. In order to reduce redundancy, all consensus sequences were clustered using a 80% identity threshold (mmseqs22 v13; -k 0 --cov-mode 1) (24). For each resulting cluster only the longest sequence was then kept in the library. To retrieve repeat positions in the genome assembly, consensus sequences from the non-redundant library were mapped to the genome using RepeatMasker3 (v4.1.2, -no_is) (25).

Tandem repeat sequences were annotated with Tandem Repeat Finder (TRF) v4.09.1 (26), using the parameters matching weight = 2, mismatch penalty = 7, indel score = 7, match probability = 80, indel probability = 10, minimum alignment score = 50, and motif size up to 2000 bp.

#### Immunostaining

Male *Timema* gonads exhibit a grape-like appearance, with the shoot containing mature sperm cells and the grape forming an oval structure encompassing cells in various meiotic stages (Figure 1A). Adult gonads were dissected in 1X PBS, fixed in a solution consisting of 2% paraformaldehyde and 0.1% Triton X-100 for 15 minutes, and then 4-5 "grapes" were gently squashed onto poly-L-lysine coated slides before being rapidly immersed in liquid nitrogen. After a 20-minute incubation in PBS, slides were subjected to blocking with 3% BSA blocking buffer for a minimum of 30 minutes. For immunostaining, slides were incubated with diluted primary antibodies (except for the already fluorescently labeled SMC3 antibody used in the following step) overnight at 4°C within a humid chamber. Detailed information for all antibodies used are provided in Supplemental table 5. Subsequently, slides underwent three 5-minute washes in PBS, followed by a 1-hour incubation with the secondary antibody, diluted at a ratio of 1:200 in

3% BSA, at room temperature (RT). Slides were then washed thrice for 5 minutes, followed by a 10-minute wash in 1X PBS. A blocking step was performed using diluted Normal Rabbit Serum (NRS 5%) for 30 minutes at RT, followed by a 10-minute wash in 1X PBS. Subsequently, slides were incubated for 1-hour at RT with the SMC3 antibody diluted at 1:100. Finally, slides were washed three times for 5 minutes in PBS and mounted in DAPI/Vectashield (50:50) media.

For the immuno-FISH protocol, we introduced a post-fixation step, involving incubation in a solution of 2% paraformaldehyde and 0.1% Triton for 15 minutes, before proceeding with the FISH protocol as detailed below (without the fixation and freezing steps).

#### Fluorescent In-Situ Hybridization

Tissues were fixed and squashed on slides as described for the immunostaining protocol. After a rapid freezing step in liquid nitrogen, cover slips were removed and slides immersed in PBS containing 0.1% Tween20 for 20 minutes. For the hybridization step, 1 µl of each probe (at a concentration of 100 ng/µl) was diluted in 20 µl of 1.1x hybridization buffer (composed of 10 µl formamide, 4 µl 50% dextran sulfate, 2 µl 20X SSC, and 4 µl ultrapure water). This probe/hybridization buffer mixture was added to the slide and covered with a cover slip. The slides with coverslips were then heat-shocked for 1 minute at 95°C, and incubated at 30°C overnight within a humid chamber. Finally, slides were subjected to three 5-minute washes in 4x SSCT [200 mL 20X SSC, 0.1% Triton, 799 mL ultrapure water] and three additional 5-minute washes in 0.1x SSC [5 mL 20X SSC, 995 mL ultrapure water] prior to mounting in DAPI/Vectashield (50:50) media.

#### Image acquisition

All acquisitions characterizing CenH3 and kinetochore phenotypes during male meiosis were performed using the Zeiss LSM 880 airyscan confocal microscope equipped with a 60x/oil immersion objective. All acquisitions were produced by the superimposition of focal planes. Post-processing, including croppin and pseudocoloring, was carried out using Fiji (27).

To detect the ultrastructural organization of chromosomes, CenH3 and tubulin signals, and to pinpoint spindle microtubule attachment points at a resolution of ~120 nm (super-resolution achieved with a 488 nm laser excitation), spatial structured illumination microscopy (3D-SIM) was performed with a 63×/1.4 Oil Plan-Apochromat objective of an Elyra 7 microscope system and the software ZENBlack (Carl Zeiss GmbH). Images were captured separately for each fluorochrome using the 561, 488, and 405 nm laser lines for excitation and appropriate emission filters (28, 29). All acquisitions were produced by the superimposition of focal planes and post-processing, including editing, cropping, and pseudocoloring, was carried out using Fiji (27).

#### Chromatin preparation

To identify CenH3 binding sequences during *Timema* male meiosis, we first performed a chromatin preparation on dissected testes immediately frozen in liquid nitrogen. Eighty-five milligrams of frozen tissues were transferred to a 2-ml Eppendorf tube, homogenized by cryogenic grinding (CryoMill; Retsch GmbH) using a specific regimen (2x 60 s, 25 Hz, resting 30 s, 5 Hz) and sequentially diluted five times with 1mL cross-linking solution composed of 50 mM Hepes (pH 7.9), 1 mM EDTA (pH 8), 0.5 mM EGTA (pH 8), 100 mM NaCl, and 1% formaldehyde. The 1 mL solutions were successively transferred to a 15-ml Falcon tube and subjected to rotation at RT for 12 minutes. The cross-linking reaction was stopped by pelleting nuclei for 2 minutes at 2000g, followed by replacement of cross-linking solution with a stop solution and rotation for 10 minutes. The stop solution contained 1x PBS, 125 mM glycine, and 0.01% Triton X-100. Nuclei were then subjected to washing steps in solution A [10 mM Hepes (pH 7.9), 10 mM EDTA (pH 8), 0.5 mM EGTA (pH 8), and 0.25% Triton X-100] and solution B [10 mM Hepes (pH 7.9), 1 mM EDTA (pH 8), 0.5 mM EGTA (pH 8), 0.01% Triton X-100, and 200 mM NaCl] during 10 minutes each at RT. Each washing step was followed by 2 minutes centrifugation at 2000g upon which the supernatant was discarded. After the second centrifugation, nuclei were suspended in 100 µl of radioimmunoprecipitation assay (RIPA) buffer [10 mM tris-HCl (pH 8), 140 mM NaCl, 1 mM EDTA (pH 8), 1% Triton X-100, 0.1% SDS, 0.1% sodium deoxycholate, and 1× cOmplete protease inhibitor cocktail] and transferred to AFA microtubes for sonication. Sonication was performed in a Covaris S220 sonicator for 5 minutes with a peak incident power of 140 W, a duty cycle of 5%, and 200 cycles per burst. The sonicated chromatin was transferred to a 1.5-ml Eppendorf tube and centrifuged at maximal speed for 10 minutes at 4°C, before aliquoting the supernatant to 10 µl input and 90 µL ChIP samples.

#### Chromatin Immunoprecipitation and sequencing

ChIP was carried out using 5 µl of CenH3 antibody against 45 µl of the ChIP sample filled up to 1 ml with RIPA solution and incubated overnight at 4°C. The next day, Protein A Dynabeads (25 µl; Thermo Fisher Scientific, 100-01D and 100-03D) were added for 3 hours at 4°C, and subsequently washed eight times during 10 minutes: once with RIPA, four times with RIPA with 500 mM NaCl, once in LiCl buffer [10 mM tris-HCl (pH 8), 250 mM LiCl, 1 mM EDTA, 0.5% IGEPAL CA-63, and 0.5% sodium deoxycholate], and twice in TE buffer [10 mM tris-HCl (pH 8) and 1 mM EDTA]. Finally, ChIP and input samples were subjected to ribonuclease digestion, proteinase K digestion and reversal of cross-links at 65°C for 6 hours, before being purified with CleanNGS magnetic beads from CleanNA (GC Biotech B.V, Netherlands). The purified ChIP and input DNAs were sent to the Lausanne Genomic Technologies Facility for ChIP-seq library preparation using the NEBNext Ultra II DNA Library Prep Kit for Illumina and sequencing on two Illumina HiSeq lanes (150-bp paired-end).

#### Centromere sequence identification

We first assessed whether specific genomic features (i.e., exons, introns, tandem repeats, DNA transposons, etc; see Figure 2B) were enriched in the ChIP data compared to the genomic background. This was done for the genome overall as well as separately for the autosomes and the X chromosome. We trimmed ChIP and input reads using trimmomatic (v0.39). Trimmed reads were then mapped to our reference genome using the BWA-MEM algorithm v0.7.17 (-c 1000000000). Chimeric reads were removed using SA:Z tags, and PCR duplicates were eliminated with Picard (v2.26.2). Mean coverage was computed for ChIP and input reads within non-overlapping 10 kb windows across all scaffolds using BEDTools (v2.30.0) and normalized by the number of mapped reads in each library. For each genome feature, we then summed the total length of all portions for which the mean ChIP coverage was at least 16x higher than the input coverage (i.e., $\log2(\text{ChIP/input}) \geq 4$). The frequency of these enriched features was then compared to the frequencies of genome features in the assembly (for the whole genome, the autosomes or the X) using chi-square tests.

We found that tandem repeats were the major genomic feature enriched in the ChIP data (see main text and Figure 2B). To characterize centromere sequences in *T. douglasi*, we therefore focused solely on tandem repeats. We used two different approaches, an assembly-based and an assembly-free approach. For the assembly-based approach, we defined enriched CenH3 windows (hereafter centromere windows) as 10 kb windows with a mean ChIP coverage at least four times higher than the mean input coverage (i.e., $\log2(\text{ChIP/input}) \geq 2$). We filtered all genomic features from the centromere windows that were not tandem repeats or that were tandem repeats but with a local coverage log2(ChIP/input) ≤ 2. We then categorized these enriched tandem repeats into centromere repeat families in two steps. First, we generated a catalog of unique sequence motifs among the tandem repeats using a custom Perl script. This script identified minimal rotations (including reverse complements) of all repeat motifs found by TRF. In the second step, we investigated the sequence similarity of unique motifs by calculating a Levenshtein distance for each pairwise comparison. To facilitate comparisons, all motif sequences were adjusted to the size of the longest sequence by tandem duplications whereby we used a custom python script to consider all rotations (i.e., all possible starting positions) of one of the two sequences and output the combination with the lowest Levenshtein distance. Pairwise comparisons of sequence motifs with at least 80% sequence similarity were further selected to build a network of sequence similarities from which distinct repeat families were defined (Supplemental figure 4A).

For the second approach to identify centromere repeat families, we employed a k-mer based analysis. We used the pipeline from (30) to construct k-mer databases for CenH3 ChIP-seq and input datasets with a k-mer length of 25 bp. A k-mer had to be found at least 100 times in the CenH3 and input datasets to be included in the k-mer database. We counted and normalized the abundance of each k-mer relative to the total base pairs. Centromere enrichment values were determined by calculating the ratio of normalized counts in the CenH3 dataset to those in the input dataset. Enriched k-mers were identified as those with a centromere enrichment score exceeding 25 median absolute deviations from the median (Supplemental figure 4B). CenH3-ChIP reads containing enriched k-mers were further assembled into *de novo* contigs using Spades v3.15.3 (-careful) (31). We then annotated tandem repeats within these *de novo* contigs using Tandem Repeat Finder, following the same parameters as those used for annotating the genome (see above). Tandem repeat sequences identified in the *de novo* contigs were categorized into repeat families following the same methodology as in the assembly-based approach. In short, we generated a catalog of unique motif sequences, calculated a Levenshtein distance for each pairwise comparison, and built a network of sequence similarities (Supplemental figure 4C).

The assembly-based and assembly-free k-mer approaches identified a highly congruent set of centromere repeat families, with the families identified via the assembly-free approach representing a subset of those identified via the assembly-based approach (Supplemental figure

4; Supplemental tables 2 and 3). We therefore retained the centromere repeat families identified via the assembly-based approach for further analyses.

To examine the representation and organization of the centromere repeat families on each scaffold (including the shorter scaffolds not anchored to the 12 chromosomes), we used two different methods, with very similar results. For the first method, we summed, per scaffold, the TRF-inferred lengths of all repeat arrays for motifs belonging to a specific repeat family within centromere windows. For the second method, we conducted a BLASTN search of the motifs grouped into repeat families in the centromere windows. Blast hits with sequence identity and alignment coverage of the query below 80% were excluded. This alignment threshold was chosen to fit the alignment threshold applied by TRF to identify tandem copies (https://tandem.bu.edu/trf/desc). BLASTN hits were then assigned back to specific repeat families based on query identity (Supplemental table 4) and visualized by extracting BLASTN hit coordinates within Geneious Prime (v2023.1.1). The distribution and frequency of the centromere repeat families among scaffolds were then visualized using heatmaps (Supplemental figures 5A, 5B).

#### Design and selection of antibodies and probes

To design custom antibodies for *Timema* centromere proteins, we first used CenH3, CenpC and Ndc80 protein sequences identified in (32) as initial queries to conduct tblastn searches against our *T. douglasi* gene annotations. For the CenH3 protein, three hits with E-values below $10^{-05}$ were identified, of which two were to the H3 and H3.3 histone units and were not considered further. The best hit corresponded to the CenH3 ortholog as revealed by Protein BLAST against the non-redundant NCBI database. For the CenpC and Ndc80 proteins, only a single hit with an E-value below $10^{-05}$ was identified. Finally, we also conducted tblastn searches for the sequences of the three proteins against our *T. douglasi* gene annotations to identify possible paralogs, but only secondary hits with low percent sequence identity and coverage were recovered. We then used the three protein sequences to outsource to Covalab (Lyon, France) the design of three distinct peptide sequences for each protein (Supplemental table 5). Follow-up tblastn searches against our *T. douglasi* gene annotations were performed to corroborate the absence of peptide cross-reactions with non-target proteins. None of the peptides had cross-reactivity with other annotated genes for the CenH3 and Ndc80 proteins, while peptide 2 from the CenpC protein had a potential but unlikely cross-reactivity with the annotated, anonymous gene Tdi_034212-RA (i.e., 46% sequence identity). Covalab subsequently developed polyclonal rabbit antibodies for the three target genes using the designed peptides, and purified them via a sepharose column.

We assessed the specificity of the CenH3 custom antibody using western blot analyses. First, nuclear enriched proteins were extracted from frozen *Timema* testes. Testes were ground for 1 minute at 25 Hrz in liquid nitrogen using Cryomill (Retsch) and resuspended in 600 µL of TEB (PBS1x; 0.5% Triton X100 (v/v); cOmplete Protease Inhibitor Cocktails (Sigma-Aldrich, #11697498001, 1 tab in 50 mL)). After a short spin at 4°C to remove debris, the supernatant was transferred to a new tube and centrifuged for 10 minutes at 4°C at 2000 rpm. The supernatant was discarded and the pellet was resuspended in 40 µL of Laemmli buffer (1x) followed by 5 minutes incubation at 95°C. Final protein concentration was measured using tryptophan fluorescence according to (33).
Second, a western blot was run from the nuclear enriched proteins and revealed a single band at the expected size for CenH3 (Supplemental figure 6). 1.5 µg of nuclear enriched proteins and 25 µL of pre-stained protein standard (Bio-Rad #161-0377) were loaded into a 4-10% precast polyacrylamide gel (Bio-Rad #456-1093) and run for 30 minutes at 180V in a Mini-trans-Blot Module following Bio-Rad recommendations. Proteins were then transferred to nitrocellulose membrane (Bio-Rad #1620112) in wet conditions for 50 minutes at 4°C and 60V according to Bio-Rad guidelines. A 5 minute ponceau (Sigma-Aldrich #P7170) staining was performed to check the quality of the transfer. A blocking step was performed using NFDM (Not Fat Dry Milk) 3% in PBT (PBS 1x + 0.1% Tween-20) for 1h at RT. The membrane was then incubated overnight at 4°C with a solution of primary CenH3 antibody (diluted 1:3000 in NFDM 3%). Three washes of 10 minutes with PBT was performed before staining with anti-Rabbit HRP secondary antibody (Jackson #111-035-144, diluted 1:5000 in NFDM 3%) for 1h at RT. After three washes in PBT, Clarity Western ECL Substrate kit (Bio-Rad # 170-5060) was used for revelation, following instructions. Chemiluminescence acquisition of the membrane was made with a Fusion imaging system (by Vilber) with 20 seconds exposition.

To corroborate the centromere localizations of the CenH3-enriched motifs identified from the ChIP-seq data, we selected four of the most represented repeat families to design FISH probes that were custom-ordered from Microsynth (Balgach, Switzerland) and labeled using a 5' modification with single fluorophore (Supplemental table 5). In designing probes, we selected a representative motif within each of the four abundant families by trying to maximize its overall abundance (in base pairs) across centromere windows, the number of occurrences, and its connectivity within the network of CenH3-enriched motifs.

### Supplemental Results

#### Supplemental tables

**Supplemental table 1.** *T. douglasi* genome assembly statistics. The size of the *T. douglasi* genome assembly is approximately 1.29 Gb, fitting well with estimates from previous short-read assemblies (14), and high BUSCO scores reveal that it is largely complete. The number of large scaffolds assembled (n=12) matches the number of chromosomes inferred from karyotypes (11 autosomes and 1 X chromosome; (11)) and comprises 98.4% of the total assembly. The annotation of the total assembly includes 39688 genes, 32.2% of TEs and 22.6% of TRs.

| Species (accession number) | Total assembly size (all scaffolds) | Size of the 12 large scaffolds | Total number of scaffolds | BUSCO v5, n:1367 |
|---|---|---|---|---|
| *T. douglasi* (PRJNA1123914) | 1.29 Gbp | 1.27 Gbp | 1136 | C:99.0%[S:98.2%, D:0.8%],F:0.7%,M:0.3%<br>4 missing. |

**Supplemental table 2**. Tandem repeat motifs assembly-based method. https://docs.google.com/spreadsheets/d/1imtiBwwIKrJ9Hs-YmXkxdGqYNwIaUJGhpR-K6LkRijc/edit#gid=1709228053

**Supplemental table 3**. Tandem repeat motifs reference-free method. https://docs.google.com/spreadsheets/d/1opS85CAZcLkN4nDgvy4_zS3a299OcVtdYKbiBEpOdj4/edit#gid=0

**Supplemental table 4**. Blast hits of tandem repeat motifs against centromere windows. https://docs.google.com/spreadsheets/d/1JaLwMPj1m-FYBaXejm9eHWD7CP0Tgtyi/edit?usp=sharing&ouid=106478781139969559754&rtpof=true&sd=true

**Supplemental table 5**. Antibodies and probes used. A combination of custom and commercially available antibodies as well as DNA probes was employed for the immunostaining and FISH assays. The CenH3-custom antibody was further used in the ChIP-seq and ImmunoFISH experiments.

| List of primary custom antibodies | | | |
|---|---|---|---|
| **Short Name** | **Finale Concentration** | **Peptide sequence** | **Tdi gene annotation reference** |
| | | VRRKSSAKKRSIRISGPREET-C-coOH | |
| | | C-ETSARSNKTQNDSSKPSTSH-coNH2 | |
| CenH3 | 1/100 | C-SKPSTSHHKSKNKSTRWSG-coNH2 | Tdi_018724-RA |
| | | C-ESSVREVTKSSSGGS-coNH2 | |
| | | C-KVHNKTKQTSKGRNKT-coNH2 | |
| CenPC | 1/100 | C-TYTKHGAEELSGSGE-coNH2 | Tdi_024923-RA |
| | | C-FSSNKGSQKNKNTYAT-coNH2 | |
| | | C-DKVGFSEAEEQKYLE-coNH2 | |
| NDC80 | 1/100 | C-KEESAQAEYKQEREK-coNH2 | Tdi_023905-RA |
| List of primary commercial antibodies | | | |
| **Short Name** | **Reference** | **Company** | **Finale Concentration** |
| SMC3 | ab201542 | AbCam | 1/100 |
| α_Tubulin | #F2168 | Sigma-Aldrich | 1/150 |
| List of secondary antibodies | | | |
| **Short Name** | **Reference** | **Company** | **Finale Concentration** |
| anti-Rabbit-Alexa 594 | 711-585-152 | Jackson | 1/150 |
| List of probes | | | |

| Short Name | 5' modification | Probe sequence |
|---|---|---|
| Repeat family 1 | Alexa fluorophore 488 | GAAGATTATTGAAATCAAGTATGTCCGCTTTGTTGATTATTTTCCG |
| Repeat family 7 | Fluorophore Cy5 | CTAAATAATCAAAATCGGCTATGGATGCTCGGTTGACAAGTTTTATC |
| Repeat family 13 | Alexa fluorophore 594 | AGCTGAAGTGTGCAAGTATTTGTGGGAGGATACCATACCCGGCGTCTAGT |
| Repeat family 22 | Alexa fluorophore 594 | TAGTTAGGCTAGATCAGCCAGTCAGGTCGTATCCCCGTATCTTATAGAGT |

### Supplemental Figures

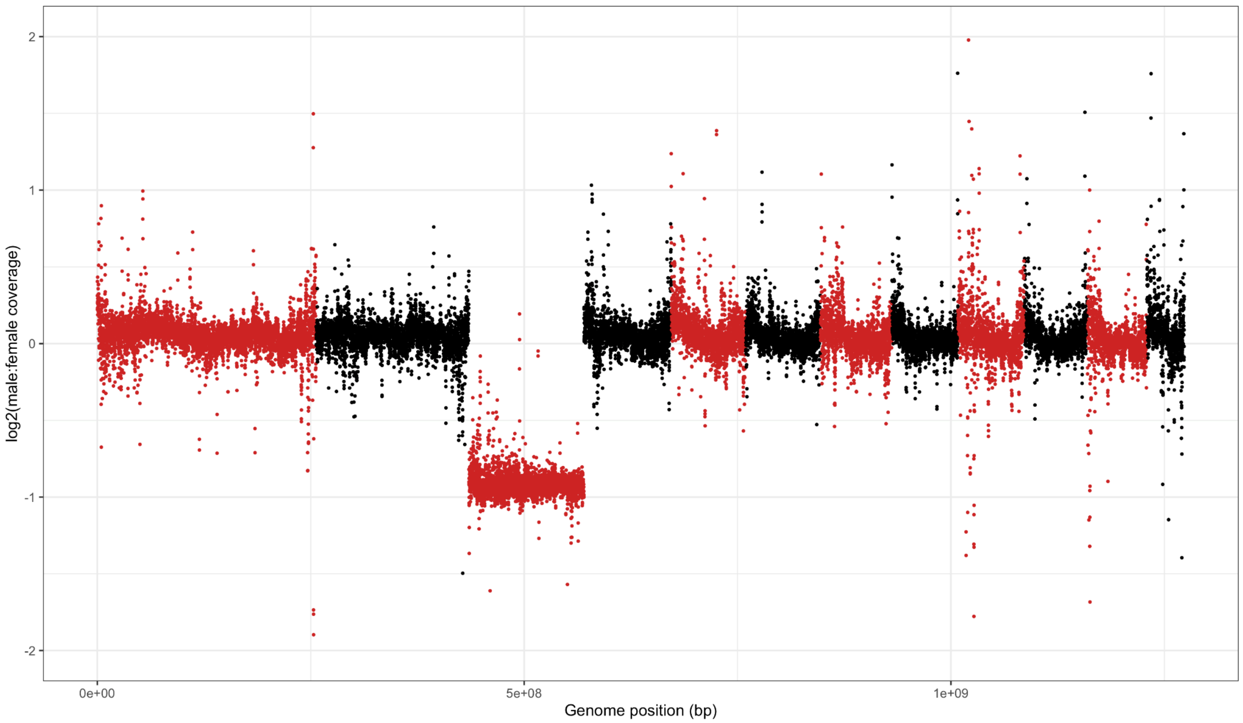

**Supplemental figure 1.** ***T. douglasi*** **X chromosome identification.** The plot shows the $log_2$ ratio of male to female coverage of 100 kb sliding windows across the genome. Alternated colors designate different chromosomes, with chromosome 3 showing a much lower overall male to female coverage ratio.

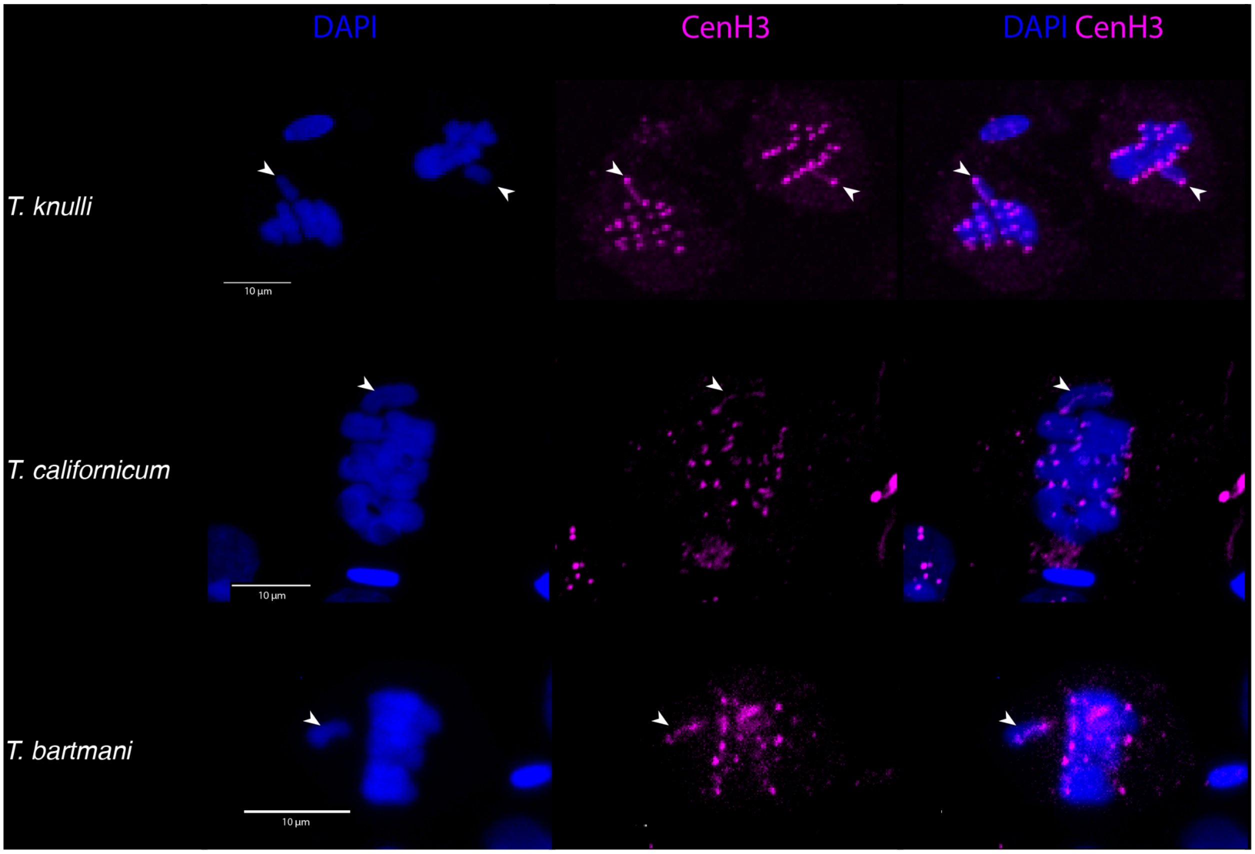

**Supplemental figure 2.** Mono- and holocentric-like CenH3 binding along autosomes and the X chromosome, respectively, for meiotic cells of males from three different *Timema* species.

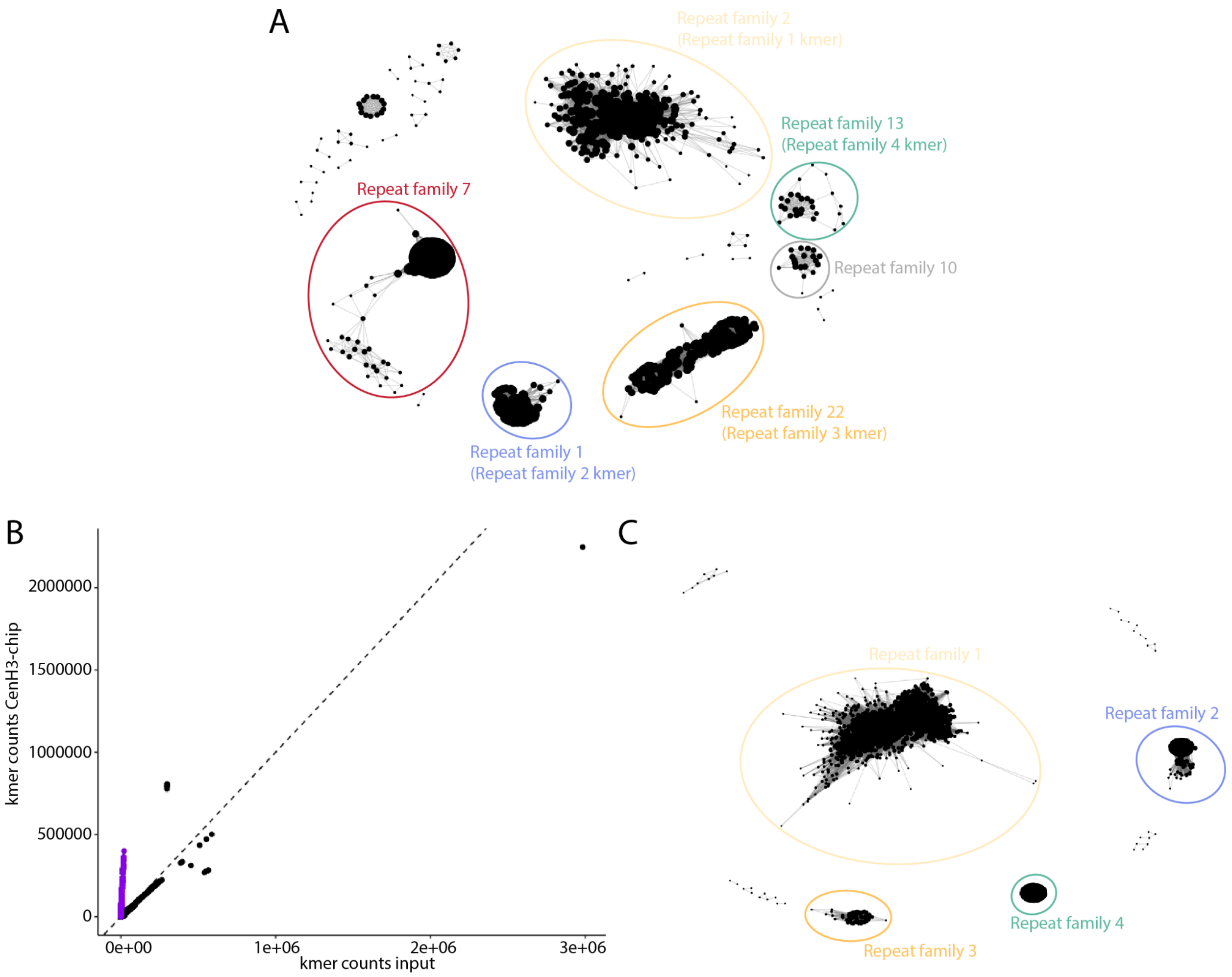

**Supplemental figure 3.** Identification of centromere sequences. A) Network of sequence similarities among tandem repeat motifs identified in the genome assembly. In the two networks, each node represents a unique CenH3-enriched motif sequence and edges connect motifs with at least 80% sequence similarities. B) Scatterplot of 25-bp k-mer normalized counts found in input and CenH3 ChIP-seq libraries. Enriched k-mers are highlighted in purple and were identified as those with a centromere enrichment score exceeding 25 absolute deviations from the median. C) Network of sequence similarities among tandem repeat motifs identified in *de novo* contigs (k-mer based approach).

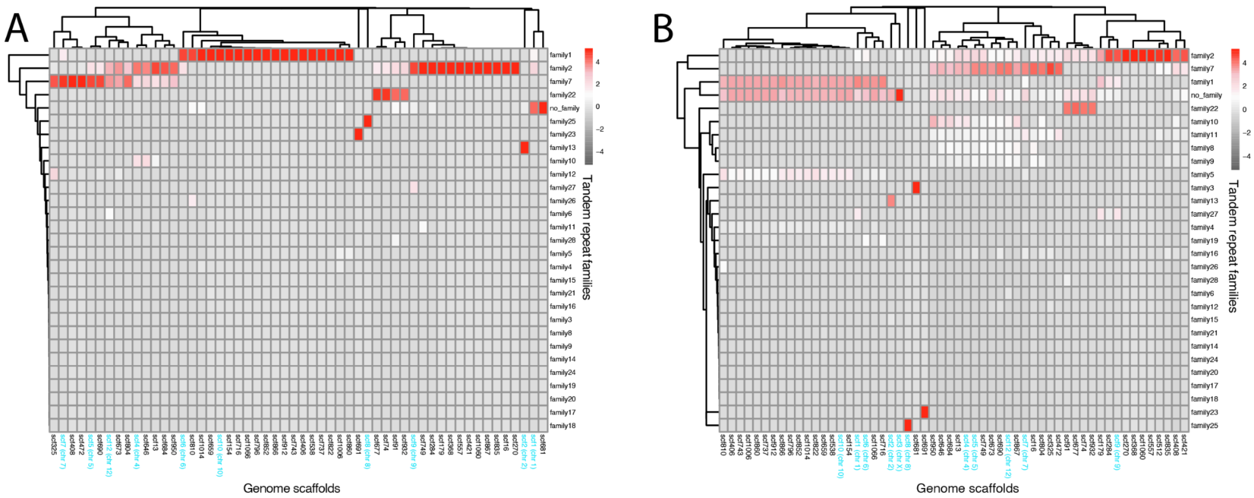

**Supplemental figure 4.** Centromeres of different chromosomes consist of distinct tandem repeat families. The heatmaps are based on the subset of scaffolds comprising centromere tandem repeat families in at least one centromere window, with hierarchical clustering based on (A) total array length inferred by summing Tandem Repeat Finder array lengths per repeat family and (B) total array length inferred by summing the lengths of sequence motif blast hits with 80% sequence similarity and 80% query coverage. Chromosome scaffolds are highlighted in cyan.

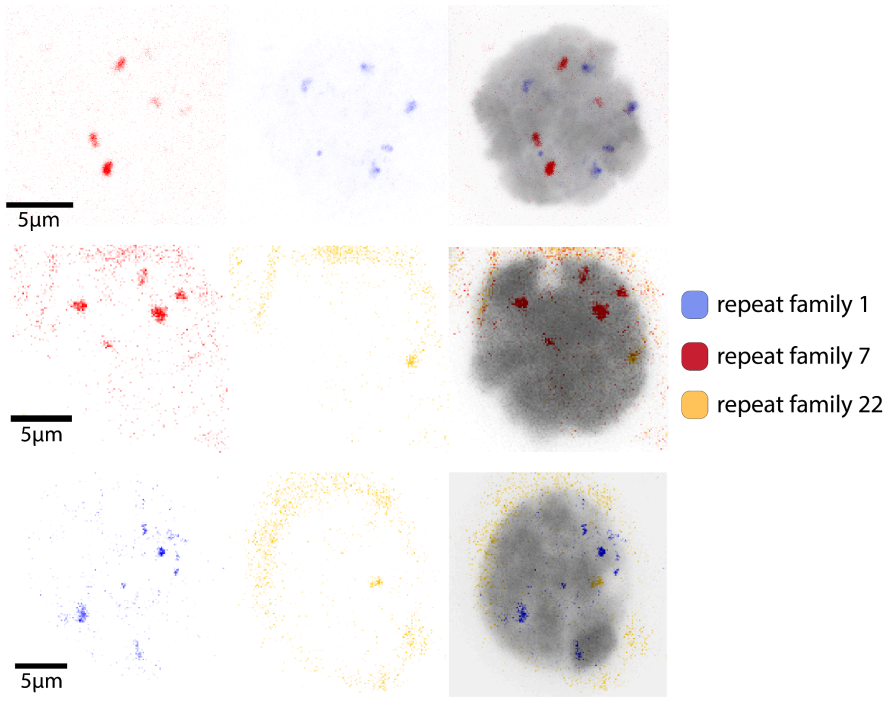

**Supplemental figure 5.** Fluorescent In Situ Hybridization of DNA probes labeling specific centromere repeat families. DNA is labeled with DAPI in grey.

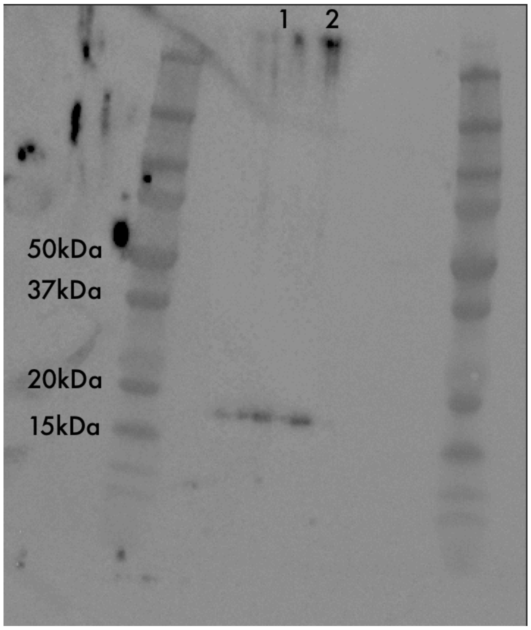

**Supplemental figure 6. Immunoblotting using the custom CenH3 antibody.** Immunoblot analysis of protein extracts from *Timema* testes, using the custom polyclonal antibody against CenH3. Molecular weight markers are indicated by numbers in kilodaltons (*left*) and their position by lines. The CenH3 polyclonal antibody is expected to recognise a band around 18 kDa in *Timema*.
